# Dilated cardiomyopathy mutation E525K in human beta-cardiac myosin stabilizes the interacting heads motif and super-relaxed state of myosin

**DOI:** 10.1101/2022.02.18.480995

**Authors:** D.V. Rasicci, P. Tiwari, R. Desetty, F.W. Sadler, S. Sivaramakrishnan, R. Craig, C.M. Yengo

## Abstract

The auto-inhibited, super-relaxed (SRX) state of cardiac myosin is thought to be crucial for regulating contraction, relaxation, and energy conservation in the heart. We used single ATP turnover experiments to demonstrate that a dilated cardiomyopathy (DCM) mutation (E525K) in human beta-cardiac myosin increases the fraction of myosin heads in the SRX state (with slow ATP turnover), especially in physiological ionic strength conditions. We also utilized FRET between a C-terminal GFP tag on the myosin tail and Cy3ATP bound to the active site of the motor domain to estimate the fraction of heads in the closed, interacting-heads motif (IHM); we found a strong correlation between the IHM and SRX state. Negative stain EM and 2D class averaging of the construct demonstrated that the E525K mutation increased the fraction of molecules adopting the IHM. Overall, our results demonstrate that the E525K DCM mutation may reduce muscle force and power by stabilizing the auto-inhibited SRX state. Our studies also provide direct evidence for a correlation between the SRX biochemical state and the IHM structural state in cardiac muscle myosin. Furthermore, the E525 residue may be implicated in crucial electrostatic interactions that modulate this conserved, auto-inhibited conformation of myosin.

**Significance Statement:** Dilated cardiomyopathy can be caused by single point mutations in cardiac muscle myosin, the motor protein that powers contraction of the myocardium. We found that the E525K DCM mutation in the cardiac myosin heavy chain stabilizes the auto-inhibited, super-relaxed state, suggesting a mechanism by which this mutation reduces muscle force and power. The E525K mutation also highlights critical electrostatic interactions important for forming the conserved, auto-inhibited conformational state of striated muscle myosins.

## Introduction

Muscle contraction is driven by the sliding of thick and thin filaments in the muscle sarcomere. In striated muscle, contraction is regulated by both thin and thick filament mechanisms. It is well-established that a rise in intracellular calcium concentration changes the conformation of the actin-containing thin filaments (thin filament regulation), allowing myosin heads in the thick filament to bind to actin and power filament sliding (Kobayashi and Solaro, 2005). Recently, thick filament regulation has garnered much attention, as myosin heads can form an auto-inhibited or super-relaxed (SRX) state with slow ATP turnover (Hooijman et al., 2011; Nag and Trivedi, 2021; Stewart et al., 2010). Structural studies have shown that myosin heads in the thick filament can fold back on the myosin tail and interact with each other in a conformation referred to as the interacting-heads motif (IHM) (Alamo et al., 2018; Woodhead et al., 2005; Zoghbi et al., 2008), which prevents them from interacting with actin. However, it is currently unclear if the SRX biochemical state and IHM structural state are directly correlated (Craig and Padron, 2022).

The SRX state is proposed to play several important roles in cardiac muscle. It appears to play a crucial role in conserving energy, as SRX heads turn over ATP about 5-10 times slower than the uninhibited disordered relaxed heads (DRX) (Hooijman et al., 2011; Toepfer et al., 2020). Cardiac muscle may also use the SRX heads as a reserve that can be recruited when peripheral metabolic demands warrant increased cardiac contractility (McNamara et al., 2015). It has been hypothesized that regulation of the SRX state in cardiac muscle underlies the Frank-Starling mechanism, in which contractile force increases as cardiac muscle is stretched (i.e. length-dependent activation) (Campbell et al., 2018; Kampourakis and Irving, 2021; Zhang et al., 2017).

Point mutations in cardiac myosin can lead to various forms of cardiomyopathy, with hypertrophic (HCM) and dilated (DCM) being the most common phenotypes (Yotti et al., 2019). HCM presents clinically as hypertrophy of the interventricular septum, resulting in decreased left ventricular diameter and pronounced relaxation deficits. These changes, coupled with myofibrillar disarray and fibrosis, typically result in a hypercontractile phenotype. Recent work has revealed that several HCM mutations in cardiac myosin can destabilize the SRX state (Adhikari et al., 2019; Anderson et al., 2018; Gollapudi et al., 2021; Sarkar et al., 2020; Vander Roest et al., 2021), which may explain the impaired relaxation and hypercontractile phenotype. DCM presents as thinning of the myocardium and a subsequent increase in left ventricular chamber volume, associated with cardiomyocyte cell death and a hypocontractile phenotype (McNally and Mestroni, 2017). An attractive hypothesis is that DCM mutations may stabilize the SRX state, which reduces the number of myosin heads available to produce contractile force and could explain the observed hypocontractility. However, the leading hypothesis is that DCM mutations reduce the intrinsic motor properties of myosin without altering the SRX state (Robert-Paganin et al., 2018; Ujfalusi et al., 2018), although few studies have examined the impact of DCM mutations on the SRX state.

The IHM, which appears to be a conserved regulatory structure in all muscle types (Alamo et al., 2018; Lee et al., 2018), has been proposed to be the structural basis of the SRX state (Craig and Padron, 2022). The motif is found in both single myosin molecules and native thick filaments. Early studies with smooth and non-muscle myosin revealed a folded tail conformation that was proposed to be auto-inhibitory (Craig et al., 1983; Onishi and Wakabayashi, 1982; Trybus et al., 1982; Trybus and Lowey, 1984). A study with smooth muscle myosin was the first to reveal the interactions between the heads that characterize the IHM (Wendt et al., 1999; Wendt et al., 2001), and later how the folded tail stabilizes the interacting heads (Burgess et al., 2007). The motif subsequently has been demonstrated in myosin II molecules across the evolutionary tree, from humans to the earliest animals (e.g. sponges) (Jung et al., 2008; Lee et al., 2018). Further biochemical studies demonstrated that the IHM is the auto-inhibited structure that predominates in the dephosphorylated off-state (Burgess et al., 2007; Cross et al., 1988; Wendt et al., 2001). Three recent cryo-EM studies of smooth muscle myosin solved the high-resolution structure of the IHM, providing crucial details about head-head and head-tail interactions that stabilize the structure (Heissler et al., 2021; Scarff et al., 2020; Yang et al., 2020). The IHM was first demonstrated in native thick filaments in tarantula muscle (Woodhead et al., 2005) and then was shown to be conserved in thick filaments throughout the animal kingdom, including vertebrates (Al-Khayat et al., 2013; Alamo et al., 2018; Zoghbi et al., 2008). The highest resolution so far (13 Å) has come from tarantula filaments (Yang et al., 2016), which showed an IHM with similar features to smooth muscle myosin (Alamo et al., 2018; Alamo et al., 2008). However, in skeletal and cardiac myosin the high-resolution structure of the IHM has not been solved.

In order to directly examine the correlation between the IHM and SRX state, a method that can detect the IHM in solution in addition to the fraction of molecules with slow ATP turnover is crucial. In the current study, we developed a method of detecting the IHM in solution by FRET, allowing us to directly correlate the fraction of myosin heads in the IHM with the fraction of myosin heads in the slow ATP turnover (SRX) state. In addition, we examined the impact of a DCM-associated mutation, E525K, on the formation of the IHM structural and SRX biochemical states. We found that the E525K mutation dramatically stabilizes the SRX state, monitored by single ATP turnover, as well as the IHM, monitored by FRET and confirmed by single molecule EM imaging. Our results suggest that stabilization of the off-state is a viable explanation for the decrease in muscle force and power in DCM. We also demonstrate that the IHM correlates well with the SRX state, providing evidence that the IHM is the structural basis of the SRX biochemical state.

## Results

### Overall approach and rationale

Our initial goal was to produce a human beta-cardiac myosin construct that would allow us to monitor formation of the SRX biochemical and IHM structural states in solution. Previous studies in smooth muscle myosin demonstrated that at least 15 heptads of the S2 region of myosin were required for phosphorylation-dependent regulation (Trybus et al., 1997). Thus, we generated human beta-cardiac heavy meromyosin (HMM) with 15 heptads of S2, a leucine zipper, and a C-terminal GFP tag (M2β 15HPZ.GFP) (Figure 1). We expressed and purified M2β 15HPZ.GFP using the C2C12 expression system (Chow et al., 2002; Wang et al., 2003), and the myosin contained endogenous mouse skeletal muscle light chains. This is similar in composition to our published studies on S1 (Rasicci et al., 2021; Swenson et al., 2017; Tang et al., 2021). We reasoned that if M2β 15HPZ.GFP forms the IHM, then we should be able to observe FRET between Cy3ATP bound to the active site and the C-terminal GFP tag, thus essentially functioning as a biosensor of the IHM or closed state. We generated a model of the open and closed states of M2β 15HPZ.GFP and found that the predicted distance between the active site of the free head and its C-terminal GFP tag is about 90-140Å, assuming the GFP tags are flexibly linked to the tail, which is close enough to participate in FRET. All other donor-acceptor pairs (blocked head to either C-terminal GFP tag, free head to the blocked head C-terminal GFP) have distances ∼130-140 Å (Figure 1). In contrast, the distance in the open state is predicted to be ≥250 Å, which is much too far to participate in FRET. Therefore, we proposed that the FRET biosensor would be able to detect the IHM formation in solution and that the FRET distance observed would be an ensemble average of all donor acceptor pairs, which are in the range of 90-140 Å apart in the IHM.

**Figure 1.**
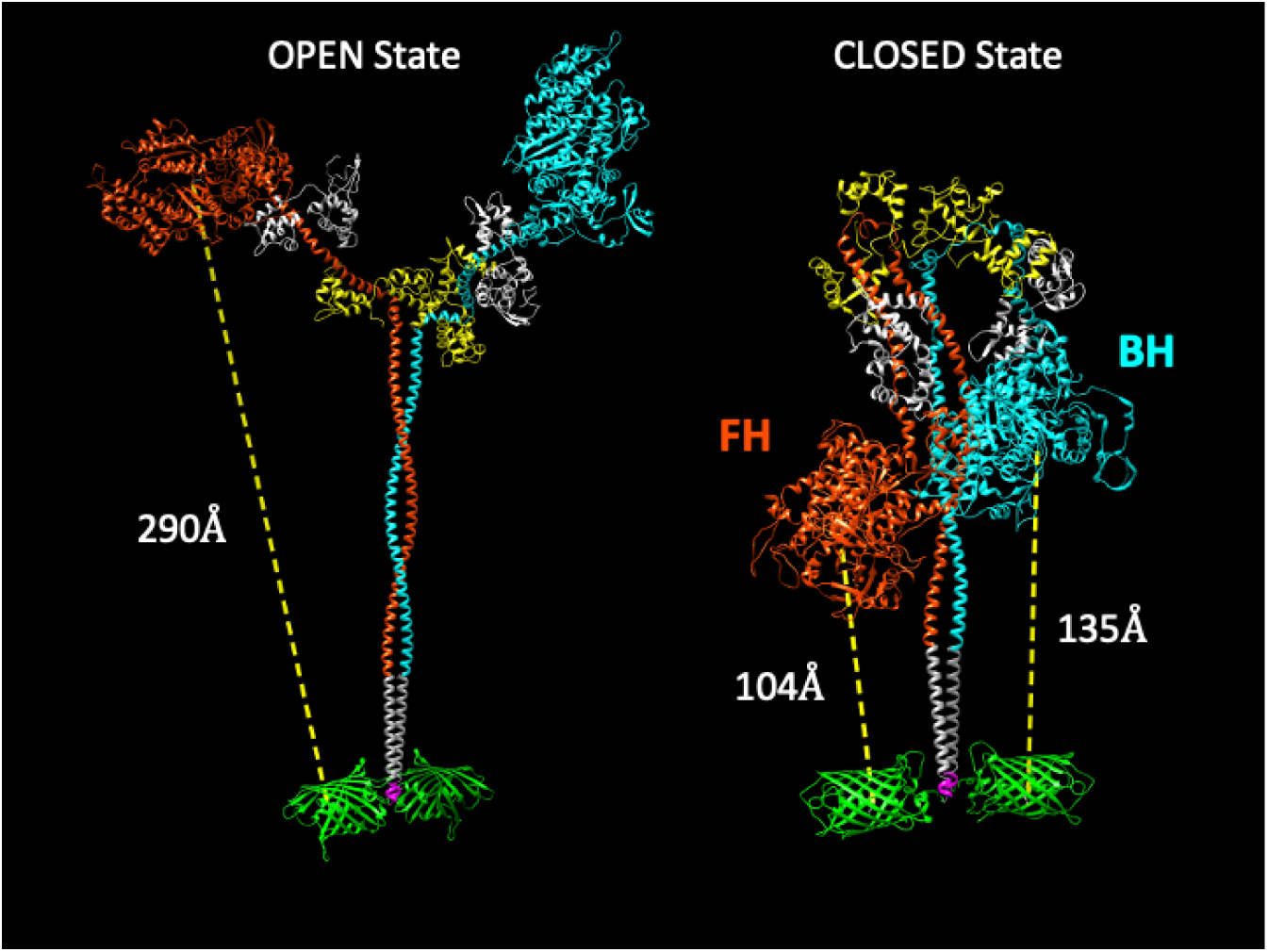
Molecular models of the M2β 15HPZ.GFP construct in open and closed (IHM) states. From proximal to distal, the model contains myosin heavy chains (orange and cyan), each with one essential light chain (white) and one regulatory light chain (yellow). Following the last residue of the 15 heptad tail (E946), the leucine zipper (PDB: 1ZIL, gray) helps to dimerize the construct, and is followed by a short, flexible glycine linker (pink) and then the C-terminal GFP molecule (PDB: 1GFL, green). Distances in the figure are measured from Y66 within the tripeptide fluorophore of GFP to S180 in the ATP binding site of the myosin motor domain. Models were constructed by joining existing pdb files in UCSF Chimera software. The open state is based on a previously published model by the Spudich lab (Trivedi et al., 2018), while the closed state is based on the 5TBY cardiac homology model from the Padrón lab (Alamo et al., 2017). In the closed state, the free head (FH) of myosin lies closer to the C-terminal GFP molecules, as compared to the blocked head (BH).

### Actin-activated ATPase and in vitro motility

In order to examine the ability of M2β 15HPZ.GFP to form the SRX state, we first examined steady-state ATPase activity in the presence of varying concentrations of actin (Figure 2A). We predicted that the ATPase activity would be decreased compared to our previous studies with monomeric S1, if the HMM construct could form the SRX state. The maximum actin-activated ATPase activity (*k*_cat_) of WT M2β 15HPZ.GFP was reduced 8-fold compared to our published measurements with WT M2β S1 in low salt conditions (20 mM KCl) (Tang et al., 2021) (Table 1). In the M2β 15HPZ.GFP construct, the *k*_cat_ was lower in the E525K mutant compared to WT while the actin concentration at which ATPase activity is one-half maximal (*K*_ATPase_) was not significantly different. We also performed *in vitro* gliding assays and found that the average *in vitro* gliding velocity was similar in WT and E525K M2β 15HPZ.GFP (Figure 2B). The velocities were similar to previous reports with a longer HMM construct (Winkelmann et al., 2015) but slower than our published results with S1 (Tang et al., 2021). There was a larger number of stuck filaments in our HMM experiments compared to S1 (Table 1), suggesting that myosin in the auto-inhibited SRX state may be able to bind actin with its free head (Wendt et al., 2001) and generate a drag force that opposes the filament sliding (see also Supplementary Movie 1).

**Table 1.** Summary of steady-state ATPase and in vitro motility measurements (±SE).

| Parameter | WT<br>M2β 15HPZ.GFP<br>(N=3-5) | E525K<br>M2β 15HPZ.GFP<br>(N=3) | #WT<br>M2β S1<br>(N=6) |
| --- | --- | --- | --- |
| $v_0$ ($s^{-1}$) | 0.02±0.02, N=5 | 0.01±0.01 | 0.02±0.02 |
| $k_{cat}$ ($s^{-1}$) | 0.99±0.37, N=5 | 0.32±0.06 | *8.34±1.12 |
| $K_{ATPase}$ (μM) | 91.2±52.6, N=5 | <sup>s</sup>36.6±14.4 | 77.5±16.2 |
| Average sliding velocity<br>(nm/s) | 853±54, N=3 | 852±58 | *1591±16 |
| Percent stuck filaments | 27.8±9.5, N=3 | 34.8±5.4 | ≤5 |

#Data is from Tang et al. 2021. \*P<0.005 comparing WT M2β S1 to WT M2β 15HPZ.GFP. <sup>s</sup>P<0.05 comparing E525K M2β 15HPZ.GFP to WT M2β 15HPZ.GFP.

**Figure 2.**
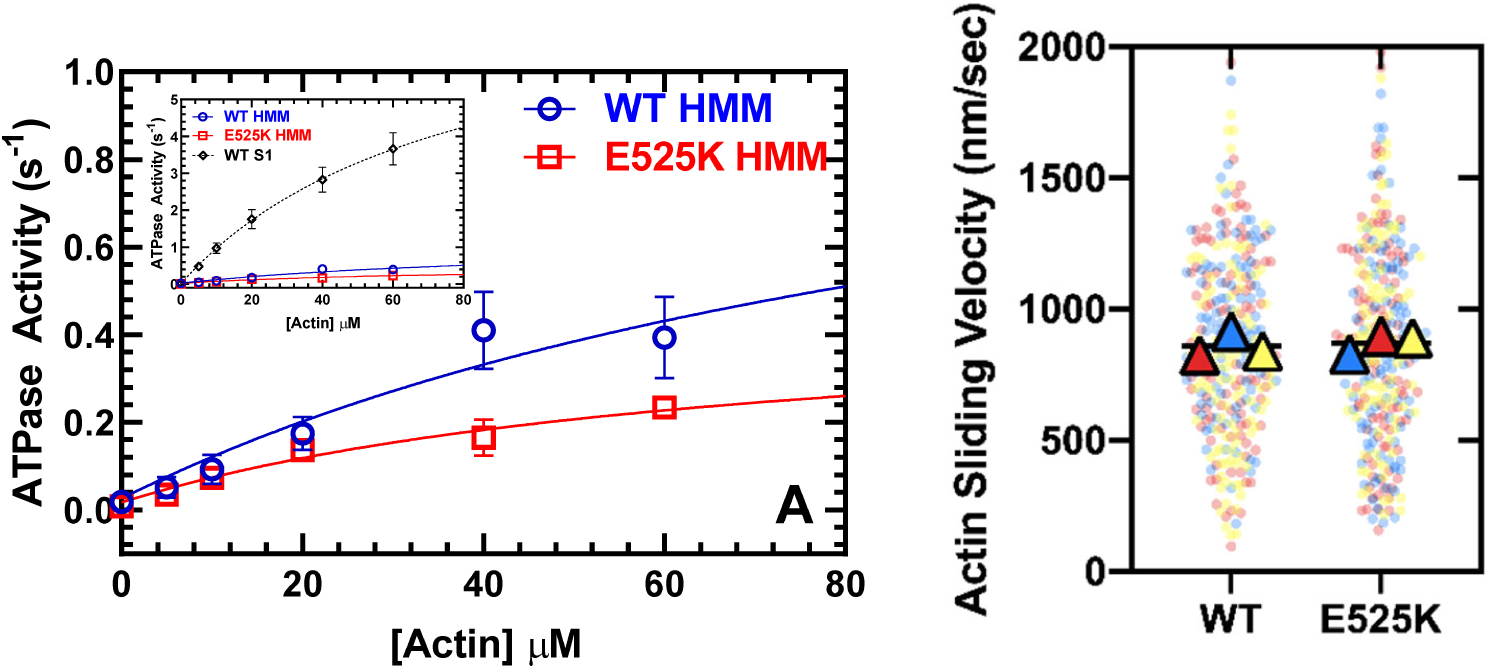
Steady-state actin-activated ATPase activity and in vitro motility. A) The ATPase activity of WT and E525K M2β 15HPZ.GFP (HMM) was measured as a function of actin concentration and the data were fit to a hyperbolic function to determine *k*_cat_ and *K*_ATPase_. Measurements are from at least 3 protein preparations (±SD). Inset demonstrates the ATPase activity of the HMM constructs in A compared to M2β-S1 (Tang et al., 2021). B) In vitro motility assays were used to examine actin gliding velocities. The individual data points are plotted for 3 protein preparations, shown in 3 different colors (yellow, blue, red), and the average for each preparation is represented by the colored triangles. The overall average velocity is represented by the black line.

### Single mantATP turnover experiments

We used mant-labeled ATP to monitor single turnover rate constants in the absence of actin (Fig. 3 and Table S1). Previous studies have demonstrated that cardiac myosin exhibits two phases in single turnover measurements, a fast phase that is similar to the basal ATPase rate of S1 and a slow phase that is 5-10 fold slower and referred to as the SRX state (Anderson et al., 2018; Hooijman et al., 2011; Rohde et al., 2018). Determining the relative amplitudes of the fast and slow phase rate constants allows determination of the fraction of myosin molecules in the open (DRX) and SRX states, respectively. We performed single turnover measurements at various KCl concentrations and observed that KCl destabilizes the SRX state in WT M2β 15HPZ.GFP, becoming a minor component at 150 mM KCl (Figs. 3A, C). However, E525K exhibited a much higher fraction of molecules in the SRX state compared to WT, especially at higher KCl concentrations (Figs. 3B, C). Thus, our results demonstrate E525K stabilizes the SRX state even under physiological ionic strength conditions. The observed rate constants for the fast and slow components of the transients were relatively similar (∼within 2-fold) in WT and E525K (Figure 3D and Table S1) suggesting the mutation mainly alters the fraction of molecules in the SRX state but does not have a major impact on the intrinsic rate of ATP catalysis in the active site.

**Figure 3.**
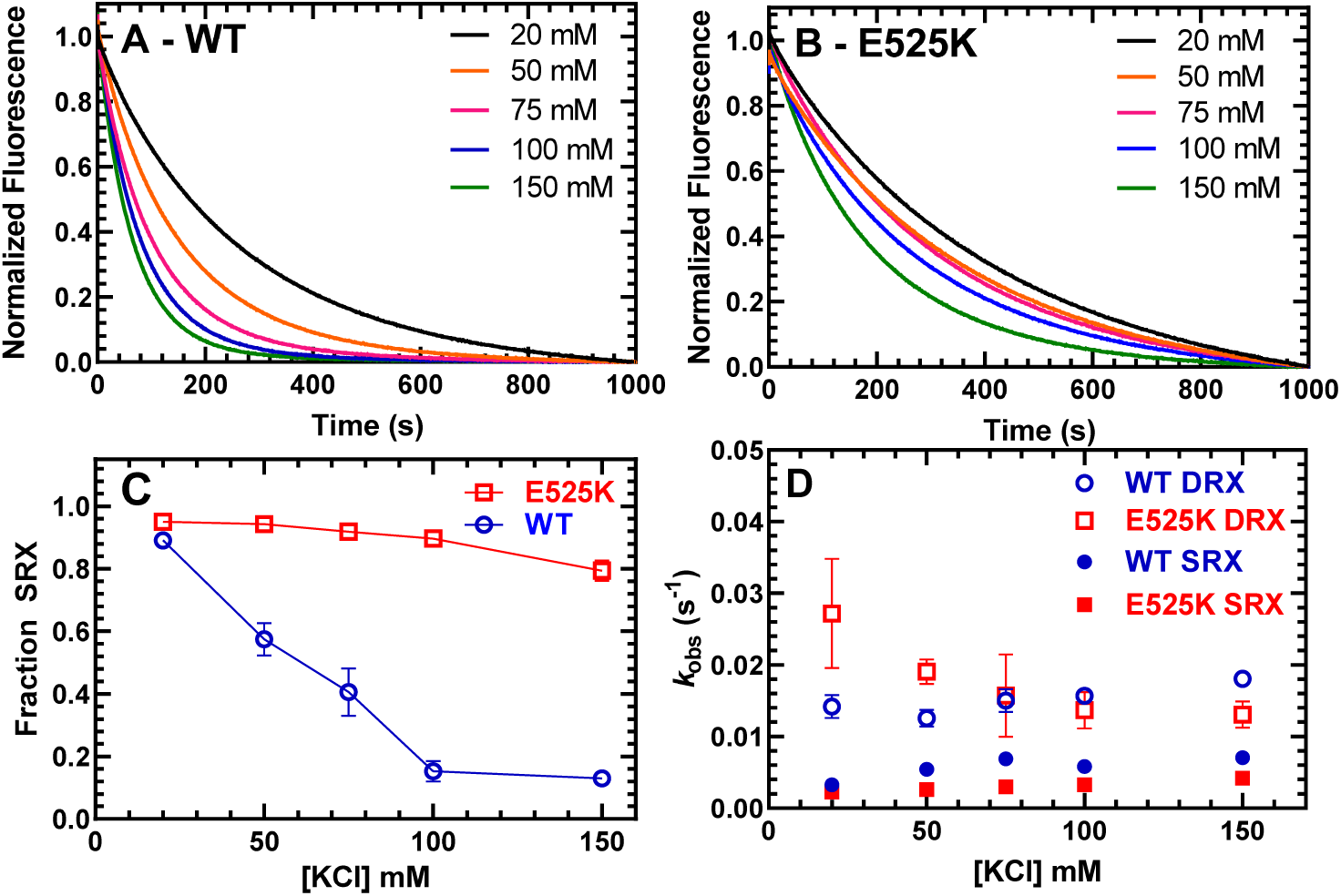
Single ATP turnover measurements. The turnover of mantATP by M2β 15HPZ.GFP was examined at varying KCl concentrations. The fluorescence transients from 0.25µM (A) WT or (B) E525K M2β 15HPZ.GFP that was pre-incubated with 1µM mantATP for ∼30 seconds and then mixed with saturating unlabeled ATP (2 mM) were observed for 1000 seconds. The mant fluorescence transients were best fit to a two-exponential function. C) The relative amplitude of the slow rate constants from the fluorescence transients were used to determine the fraction of heads in the SRX state. D) The slow rate constants (SRX state) were 5-10 times slower than the fast rate constants (DRX state) and relatively similar (∼within 2-fold) at each KCl concentration for both the WT and E525K constructs. Error bars are ±SD, N=3 (see Table S1 for summary of values and statistical comparisons).

We also monitored the mantATP single turnover rate constants in WT and E525K M2β S1. We found that the slow phase that represents the SRX state was a small fraction of the fluorescence transients (∼5%), similar in WT and E525K, and independent of ionic strength (Figure S1). The rate constants of the SRX and DRX states were 2-4 fold faster in E525K compared to WT M2β S1 (Table S2).

### Cy3ATP binding to M2β 15HPZ.GFP monitored by FRET

We tested the IHM FRET biosensor by mixing Cy3ATP with M2β 15HPZ.GFP in a stopped-flow apparatus and monitoring the quenching of the GFP fluorescence as a function of time. The approach allowed us to measure the degree of donor quenching (amplitude) as well as the rate constants for Cy3ATP binding (*k*_obs_) to M2β 15HPZ.GFP. The FRET transients clearly demonstrated donor quenching in the presence of the acceptor while donor only controls exhibited no change in fluorescence (Figure 4A&B). We reasoned that if more myosin heads were in the IHM conformation we would observe a higher FRET efficiency. The fluorescence transients fit well to a two-exponential function at low salt (20-75 mM KCl) and mostly a single exponential function at high salt (100-150 mM KCl) (Figure 4A&B). We plotted the total amplitude of the FRET change upon Cy3ATP binding as a function of Cy3ATP concentration and found that WT had a smaller amplitude at high salt (150 mM) compared to low salt (20 mM) (Figure 4C). However, the total amplitudes with the E525K experiments were quite insensitive to KCl concentration (Figure 4D), suggesting more M2β 15HPZ.GFP molecules were in the IHM conformation at higher salt. All of the rate constants were linearly dependent on Cy3ATP concentration (Fig. 4E, F, & Table S3), suggesting they represent second-order rate constants for ATP binding. The relative amplitude of the fast phase was higher in WT (65%) than E525K (25%) at low salt (Figure 4A, B & Table S3). Overall, the IHM FRET experiments followed a similar trend compared to the single ATP turnover experiments, suggesting the SRX biochemical state correlates with the IHM structural state.

**Figure 4.**
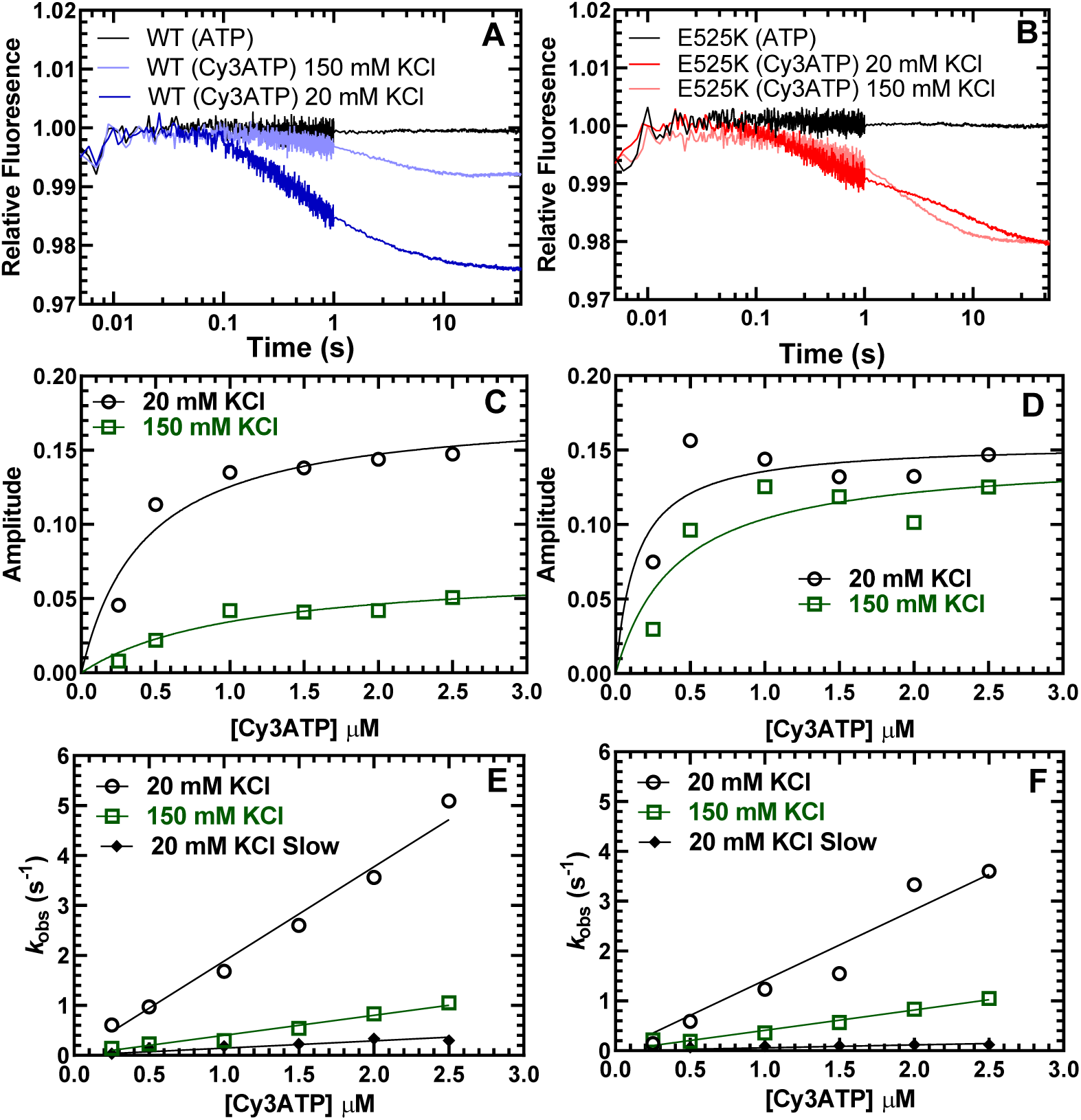
Cy3ATP binding to M2β 15HPZ.GFP monitored by FRET. GFP fluorescence (donor) quenching was examined upon Cy3ATP (acceptor) binding to WT (A) or E525K (B) M2β 15HPZ.GFP. The fluorescence transients were best fit to a double-exponential function at low salt (20 mM KCl) and a single or double exponential function at high salt (150 mM KCl) (fast phase was 65% and 25% of the signal at low salt in WT and E525K, respectively). The total amplitude of the fluorescence change was plotted as a function of Cy3ATP concentration in WT (C) and E525K (D). All rate constants were linearly dependent on Cy3ATP concentration in both WT (E) and E525K (F) M2β 15HPZ.GFP (see Table S3 for summary).

### Steady-state and time-resolved FRET measurements

The stopped-flow fluorescence transients were also used to determine the FRET efficiency after performing donor only controls. We measured the FRET efficiency as a function of KCl concentration and found that WT shifts to a lower FRET state at higher KCl concentrations while E525K maintains a similar FRET efficiency at low and high salt (Fig. 5A and Table S4). We used the steady-state FRET efficiency measurements to determine the average distance between the donor and acceptor fluorophores. This allowed us to compare the measured average distances to expected distances assuming a simplified model in which the fraction of molecules in the closed and open states directly correlates with the single turnover measurements (Figure 5B and Table S4). We assumed no FRET in the open state and a FRET efficiency of ∼5.5% associated with a 100 Å distance when all heads are in the IHM configuration. The expected distances were similar to the measured distances, especially in the E525K mutant, while the largest discrepancies were observed with WT in higher salt conditions. This is likely due to the greater degree of uncertainty with the measurements at high salt, which contained low FRET efficiency.

**Figure 5.**
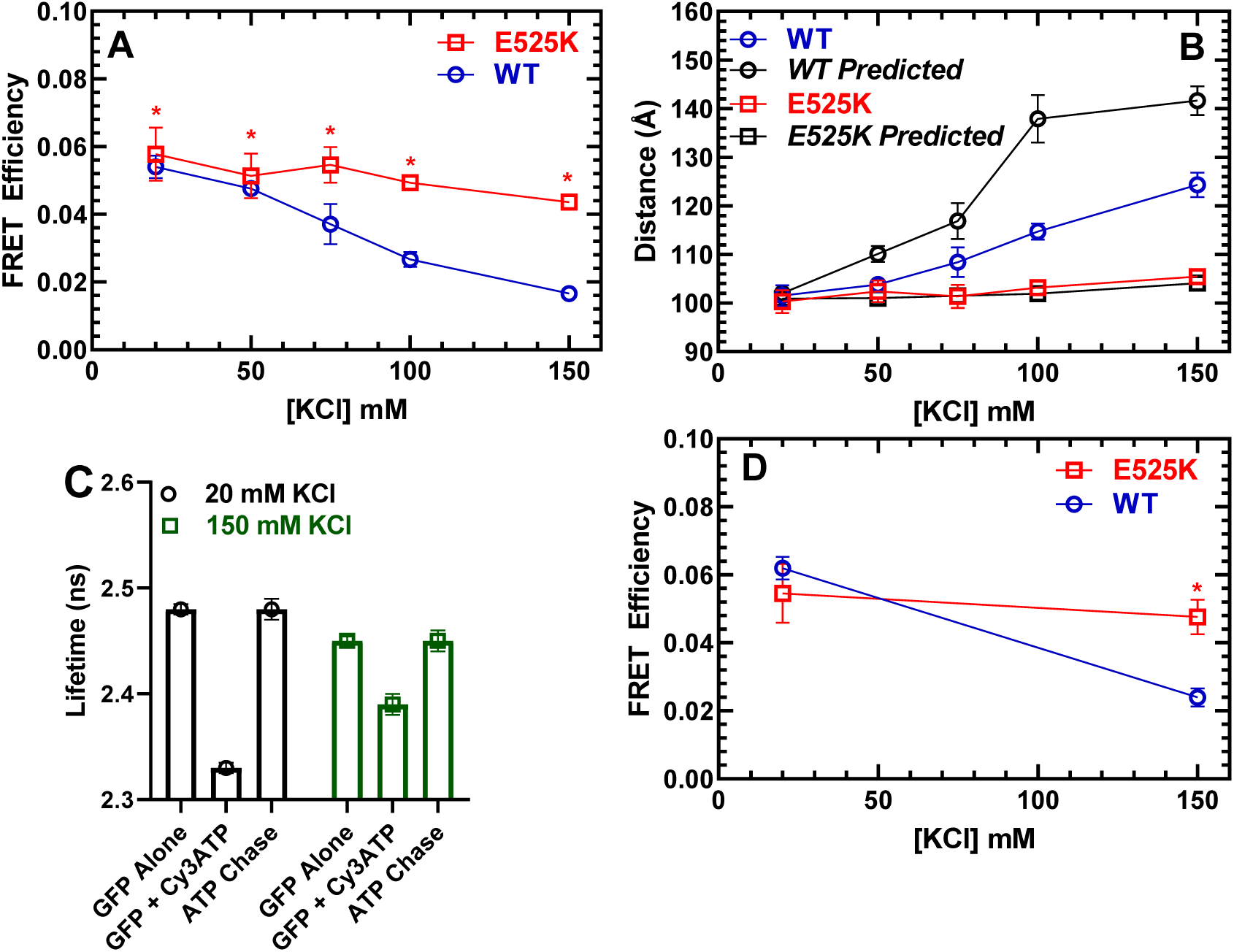
FRET monitors stability of the closed (IHM) conformation. Steady-state FRET between the C-terminal GFP tag and Cy3ATP bound to the motor domain was examined in the stopped-flow as a function of KCl concentration. A) The average FRET efficiency (±SD) was examined in 3 separate protein preparations. (B) The average distance (±SD) calculated from the FRET efficiency in panel A was compared to the predicted distance, based on a two-state model derived from the single turnover measurements in Figure 3. C) Time-resolved FRET was examined by monitoring the GFP fluorescence lifetime in the presence and absence of Cy3ATP. Representative fluorescence lifetime data with WT M2β 15HPZ.GFP. Adding saturating ATP to the sample containing WT M2β 15HPZ.GFP and Cy3ATP (ATP chase) was used to rule out donor fluorescence changes in the presence of ATP. D) Time-resolved FRET (3 technical replicates performed with a single protein preparation) demonstrated similar results to steady-state FRET shown in panel A. Error bars are ±SD, N=3 (see Tables S4&5 for summary of values and statistical comparisons) (*P<0.005).

We also performed time-resolved FRET to further verify the differences we observed between WT and E525K (Figure 5C&D). Time-resolved FRET relies on monitoring the donor fluorescence lifetime decay in the presence and absence of acceptor and thus is less sensitive to sample-to-sample variability in fluorophore concentration. We examined the GFP fluorescence lifetime (60 nM M2β 15HPZ.GFP) in the absence of nucleotide and then again after adding 0.1µM Cy3ATP. We then added saturating unlabeled ATP (1 mM) to chase off the Cy3ATP as a control for changes in donor fluorescence in the presence of ATP. The fluorescence decays were fit well to a two-exponential function which was used to determine the average lifetime in each condition. The donor fluorescence lifetimes were unaffected by the presence of ATP. We did not observe consistent differences in the relative amplitudes of the two-exponential decays (70% short lifetime / 30% long lifetime) of the GFP fluorescence that would have allowed us to perform distance distribution analysis. Overall, we found that the time-resolved and steady-state FRET measurements resulted in very similar FRET efficiencies an distances (Figure 5A&D, Table S5).

### Negative stain EM

WT and E525K M2β 15HPZ.GFP constructs were examined by negative stain EM to directly observe the impact of the mutation on the configuration of the myosin heads and to test our interpretation of the FRET data. Because the IHM is easily disrupted by binding to the grid surface during specimen preparation (Burgess 2007), we examined the molecules in low salt (20 mM KCl) conditions, which strengthen the IHM, and minimized the time of contact of protein with the grid before staining. WT molecules were dominated by non-interacting heads, producing an ‘open’ conformation (Fig. 6A, yellow circles; Fig. 6D), with only a small number (15%) of ‘closed’ (folded) IHM structures (Fig. 6A, green circles; Fig. 6D). In contrast, the majority (57%) of E525K molecules adopted the folded (IHM) conformation (Fig. 6B, green circles; Fig. 6D). To validate our assessment of open and closed percentages based on the raw micrographs (Figs. 6A, B, D), we performed 2D class averaging of the molecules using Relion 3.1 (Scheres, 2012). The class averages clearly show that in the case of E525K, most of the molecules form the IHM conformation when compared with WT, where most are open (Fig. S3). We also used class averaging to confirm that the closed structures we observed were actually IHMs. The E525K class average shown in Fig. 6C reveals key features of the IHM, including the blocked and free heads (BH and FH), the two light chains (ELCs and RLCs) and sub-fragment 2 interacting with the BH, all well-documented features of the IHM (see Figure S3). We conclude that the raw EM images and the 2D class averages support our ATP turnover and FRET results showing that E525K promotes the IHM and thus stabilizes the SRX state.

**Figure 6.**
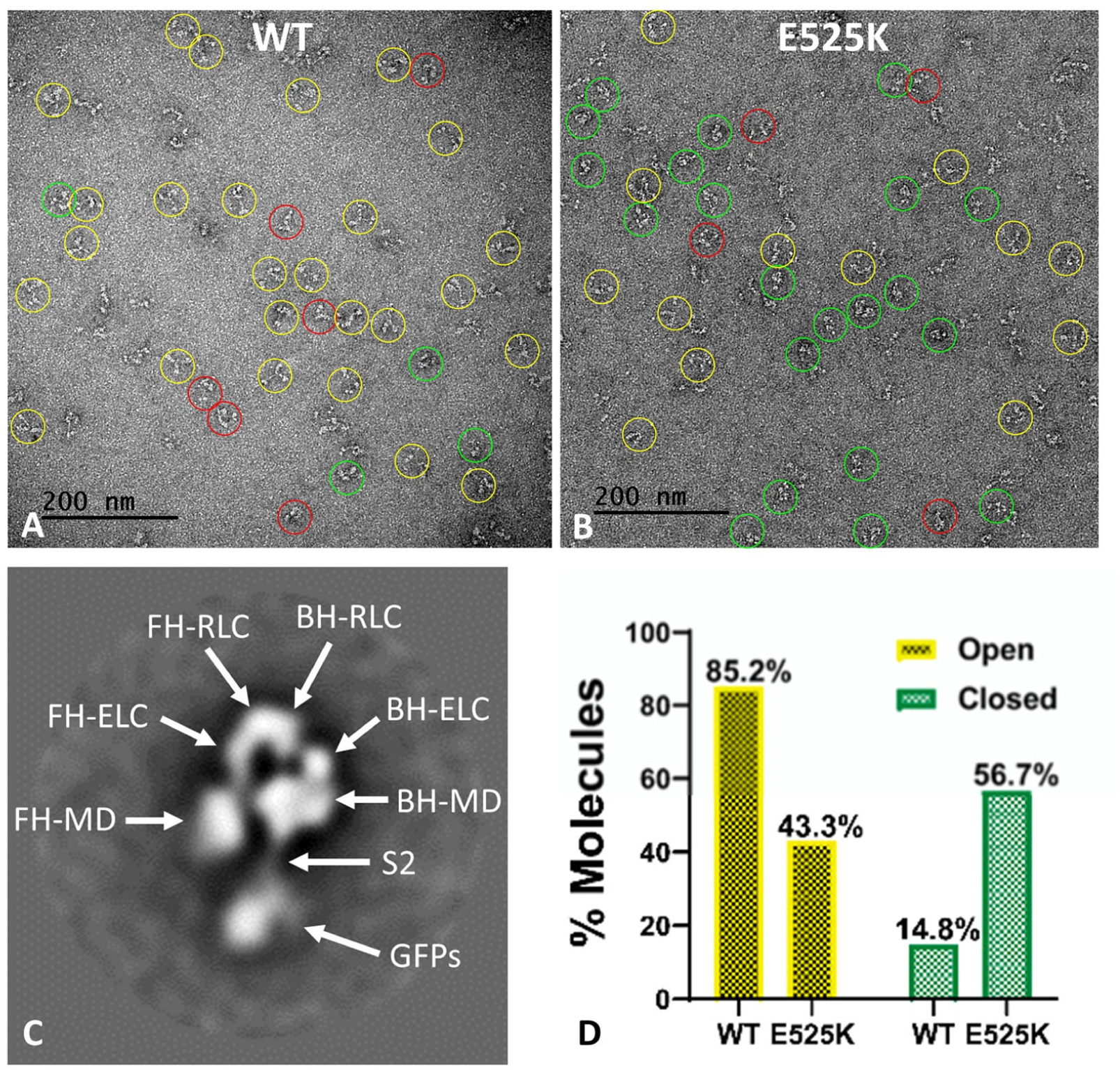
EM structure of WT and E525K constructs. A) and B) Negative stain images of WT and E525K, respectively, showing the conformational state of the molecules (yellow – open; green – closed; red circle – not clear/partially folded). C) 2D class average of E525K, showing subdomains (*cf*. Fig. 1, ‘closed’). D) Histogram showing % open *vs*. closed molecules in WT and E525K micrographs.

## Discussion

A common mechanism of myosin II regulation is via folding of the heads back onto the tail and interaction between the motor domains, effectively inhibiting actomyosin interactions and myosin ATP turnover. In the current study, we investigated the regulatory mechanism in cardiac muscle myosin, which exists in dynamic equilibrium between the folded-back conformation (IHM) and an open conformation capable of generating force upon muscle activation (Figure 1). A major question in the field is whether the IHM correlates with the SRX state, a biochemical state that turns over ATP 5-10 fold slower than single-headed S1. This proposed mechanism is attractive since the sequestered heads would inhibit interactions with actin, serving as a reserve to be recruited when enhanced contractile force is needed (e.g. length-dependent activation). Furthermore, sequestered heads would consume far less ATP, helping to balance the energetic demands of the heart. We present direct evidence that the fraction of myosin heads in the IHM correlates well with the fraction of myosin heads in the SRX state, providing crucial information about the structural basis of the auto-inhibited state of cardiac myosin. We also demonstrate that a dimeric cardiac myosin molecule with 15 heptads of the coiled-coil displays the SRX state and IHM structure, allowing us to examine the folded conformation with a novel IHM FRET biosensor. In addition, we report that a single point mutation associated with DCM (E525K) can dramatically stabilize both the IHM conformation and SRX state. Our work highlights stabilization of the IHM by electrostatic interactions, and the critical function of the E525 residue in the core region where electrostatic interactions between the head and tail occur. The clinical significance of this study is that DCM mutations can stabilize the auto-inhibited state of myosin, providing a mechanistic basis for the reduced muscle force and power observed in DCM patients.

### Correlating the SRX and IHM

The current study provides new information about the structural basis of the slow ATP turnover (SRX) state. Previously, Anderson et al. (2018) demonstrated a strong correlation between the IHM and SRX state using negative stain EM and single ATP turnover measurements. They also examined the drug mavacamten, which dramatically stabilizes the SRX state. Based on visual inspection of negative stained EM images, they observed a corresponding increase in the fraction of molecules that adopt the IHM in the presence of the drug. Our study used single turnover measurements, in addition to an IHM FRET biosensor, to examine the conformation of cardiac myosin in solution. The fraction of myosin heads in the SRX state determined by single turnover measurements was used to simulate the FRET results expected if the SRX state and IHM are 1 to 1 correlated (Figure 5B). We generated distance estimates based on the structural model shown in Figure 1 which predicts the average distance between donor-acceptor pairs in the IHM is in the range of 90-140 Å and no FRET in the open state. The measured FRET efficiency (5.5%) observed in low salt conditions corresponded to a distance of 100 Å (Figure 5A & Table S4), a condition with nearly 90% SRX heads (Figure 3C & Table S1), which allowed us to assume an average distance of 100 Å in the IHM for our simulations. The simulated data follows fairly closely with the measured FRET distances for the mutant but diverges for WT at higher salt (≥100 mM KCl) (Figure 5B), likely because the fraction of myosin heads in the IHM is quite low (≤10%) in high salt conditions and difficult to measure by FRET. Future experiments using time-resolved FRET approaches such as TR(2)-FRET (Gunther et al., 2019; Rohde et al., 2018) may allow us to determine distance distributions, distances associated with partially folded intermediates, and the rate constants for transitioning into and out of the IHM. Another study used a different FRET pair to detect the IHM and single turnover measurements with bovine cardiac myosin and found little correlation between the SRX and IHM (Chu et al., 2021). However, the FRET approach used to examine the IHM was not sensitive to ionic strength, which is very different from our results.

In addition to using FRET to monitor IHM formation, we examined negative stain EM images of the construct, both by direct visual inspection and 2D class averaging, which allowed us to systematically classify the fraction of molecules in the IHM. We found that the E525K mutant had a much higher fraction of myosin heads in the IHM compared to WT. However, this fraction was significantly different in the EM images compared to the predictions from the FRET and single turnover measurements in solution. We believe this is due to molecular interactions with the EM grid surface that destabilize the IHM. This has been well documented with smooth muscle HMM (Burgess et al., 2007). A similar but more extreme finding with cardiac myosin is exactly as predicted, given the lower stability of the cardiac compared with the smooth muscle myosin IHM (Jung et al., 2008). In fact, we found it essential to reduce the time of contact of the cardiac construct with the grid surface to a minimum (∼5 sec) before adding the uranyl acetate stain, which then immediately fixes the structure (Zhao and Craig, 2003). With longer contact times, almost all IHMs were lost, consistent with the smooth muscle HMM findings (Burgess et al., 2007). Indeed, a recent study that performed SAXS on a cardiac myosin 25 heptad construct found that alternative conformations are likely in solution, including partially folded intermediates (e.g. one head folded and one head free) (Gollapudi et al., 2021). We observed many structures by 2D class averaging with different orientations of heads. However, it is difficult to distinguish between novel conformations and states that simply lie on the grid in different orientations. Overall, our work clearly demonstrates by both EM and FRET that the IHM is a major contributor to the SRX state, while our EM studies suggest that the IHM is quite dynamic and sensitive to surface interactions.

### Structural mechanism of IHM and impact of E525K

Presently, there is no high-resolution structure of the cardiac myosin IHM. The work of Padrón and colleagues has created a model based on the tarantula thick filaments (5TBY), but using the human cardiac myosin sequence, which has been extremely useful for mapping potential head-head and head-tail interactions (Alamo et al., 2017). The recent cryo-EM structures of smooth muscle myosin in the 10S conformation provide more models of the IHM, which have some similarities and some differences compared to 5TBY (Heissler et al., 2021; Scarff et al., 2020; Yang et al., 2020).

The E525K mutation was identified in a generic study as a de novo variant of high clinical significance (Lakdawala et al., 2012). Previous structural analysis of cardiac myosin further implicates the mutation due to its location on a flat, broad region of myosin known as the mesa (Spudich, 2015). The E525 residue is located in the central 50kDa region of the myosin motor domain in Helix Q, just distal to the relay helix (Colegrave and Peckham, 2014). From the perspective of the myosin mesa, E525 is in the central region of the “mesa trail,” a region of the mesa shown to interact directly with S2 in formation of the IHM (Nag et al., 2017; Woodhead and Craig, 2020). Electrostatic attraction between the mesa trail of the blocked head (BH) and S2 is believed to underlie a crucial BH/S2 priming interaction in formation of the IHM (Alamo et al., 2017). Previous structural analyses indicate that the central mesa trail (rich in positively charged residues) interacts with the Ring 1 region of S2 (rich in negatively charged residues), and these complementary charged patches likely encourage formation of the IHM. Disruption of the charge distribution in the BH/S2 interaction region could cause complete elimination of the IHM, promoting myosin heads to populate the open or force-generating state, as has been proposed with many HCM mutations (Sarkar et al., 2020). Thus, E525 is a negatively charged residue within a positively charged mesa trail and may function normally to repel Ring 1/mesa interactions, helping maintain the delicate balance of attractive/repulsive forces required for precise functioning of the IHM. A charge-reversal point mutation such as E525K that increases the positive surface charge of the mesa trail likely would strengthen the Ring 1/mesa interaction in the BH, thus promoting the formation of the IHM. Importantly, E525 as well as many other charged residues in the mesa trail and Ring 1 of S2, is well-conserved across species and myosin isoforms, suggesting a critical role for electrostatic interactions between BH/S2 in the formation of the auto-inhibitory state (Woodhead and Craig, 2020). Without a high-resolution structure of the cardiac muscle myosin IHM, specific side chain residues in the BH/S2 interaction have not been confirmed. However, our collective data presented here coupled with previous conservation analyses indicate that E525 may be a critical residue in forming the IHM. This is further supported by our molecular modeling of the IHM, in which the introduced lysine residue at 525 of the blocked head is within several Angstroms of several acidic residues in the S2 region (Figure S4).

An alternative mechanism for how a mutation can enhance the formation of the IHM is by favoring a conformation of an intermediate that precedes IHM formation (e.g. allosteric mechanism). There is little information about the allosteric mechanisms that mediate IHM formation, while several studies have proposed that the pre-power stroke conformation of the motor domain may favor IHM formation (Alamo et al., 2017; Robert-Paganin et al., 2018). Previous studies have demonstrated that monomeric cardiac myosin (S1) can still populate the SRX state (Anderson et al., 2018; Rohde et al., 2018). Our single turnover measurements with E525K and WT S1 suggest that they have a similar ability to populate the SRX state in S1 (∼5%) and this small fraction of SRX formation is independent of ionic strength (Figure S1). Therefore, we conclude that E525K enhances the IHM and SRX state via electrostatic interactions.

### Implications for understanding the underlying mechanisms of DCM

Determining mutation-associated alterations in cardiac muscle myosin structure and function can lead to novel therapeutic strategies for the treatment of inherited cardiomyopathies such as HCM and DCM. A leading hypothesis is that HCM mutations enhance myosin-associated force and power, while DCM mutations essentially do the opposite (Debold et al., 2007; Spudich, 2014). So far, mutations in myosin associated with HCM and DCM have been found to have diverse effects on myosin motor function, including altering ATPase activity, actin sliding velocity, duty ratio, and load dependence (Trivedi et al., 2020; Ujfalusi et al., 2018). However, HCM mutations have been shown to cluster preferentially in the head-head and head-tail interaction interfaces, likely weakening interactions that underlie the SRX state, leading to greater availability of heads for actin interaction and thus to hypercontractility (Trivedi et al., 2018). The current study demonstrates that a DCM mutation can do just the opposite, dramatically stabilizing the SRX state, suggesting the possibility that other mutations may have a similar effect. A decrease in the number of available myosin heads in the thick filament would have a profound impact on contractile force and fit well with the hypocontractility hypothesis. Thus, therapeutic strategies that can destabilize the IHM and SRX state would be a logical strategy for treating DCM. However, unlike HCM, which primarily impacts sarcomeric genes, DCM is more heterogeneous and associated with a variety of genes (Yotti et al., 2019). Nevertheless, treating DCM with an SRX state destabilizer may be feasible regardless of the underlying genetic cause since it could accomplish the overall goal of enhancing contractility. The E525K mutant provides great insight into an energy conservation mechanism that may underlie development of the DCM phenotype in this patient population. Moreover, our data indicate that it can serve as a model mutation to deepen our understanding of the SRX biochemical state and IHM structural state in beta-cardiac myosin.

It is important to point out some of the limitations and alternative interpretations of the results in the current study. The M2β 15HPZ.GFP was examined with mouse light chains intrinsic to the C2C12 myocyte expression system, which could alter the impact of the E525K mutations. It has been suggested that full length myosin in a thick filament environment can further stabilize the SRX state, through interactions of IHMs with each other along the helical tracks of heads and with other thick filament proteins (Craig and Padron, 2022). Interestingly, mutations measured in purified HMM qualitatively had the same impact on the SRX state in a thick filament preparation (Gollapudi et al. 2021). The FRET approach is a biosensor of the IHM structural state while the open state is not visible by FRET, preventing us from directly determining the mole fraction of the IHM and open states. Finally, the EM provides a snapshot of a fraction of myosin molecules while the solution measurements (single turnover and FRET) provide information on the entire ensemble. The EM studies also have the limitation that the charge of the EM grid can influence the stability of the IHM.

### Summary and future directions

In summary, we examined a human cardiac myosin construct with 15 heptads in its coiled-coil and used EM imaging to show that it can form the conserved, auto-inhibited conformation known as the IHM. We also used a novel IHM FRET sensor to monitor the formation of the IHM in solution and determined that it correlates well with the SRX state with slow ATP turnover. Furthermore, we discovered that a DCM charge reversal mutation (E525K) in the motor-tail interface can greatly enhance the stability of the IHM conformation and SRX state. We conclude that this elegant method of regulation, which essentially sequesters myosin heads to balance the mechanical and energetic demands of the heart, is highly sensitive to modifications that alter the electrostatic interactions important for stabilizing the auto-inhibited conformation. Thus, future studies will investigate other DCM mutations that may also stabilize the IHM and SRX state as well as determine how other physiological factors and disease states alter this important regulatory mechanism.

## Materials and Methods

### Reagents

Cy3ATP was purchased from Jena Bioscience (1 mM stock). The 2’-deoxy-ATP labeled with N-Methylanthraniloyl at the 3’-ribose position (mantATP) was also purchased from Jena Biosciences (10 mM stock). ATP and ADP were prepared from powder (MilliporeSigma) and concentrations were determined by absorbance at 259 nm (ε_259_ = 15,400 M^-1^cm^-1^). All assays were performed in MOPS 20 buffer (10 mM MOPS, pH 7.0, 20 mM KCl, 1 mM MgCl_2_, 1 mM EGTA, and 1 mM DTT).

### Protein expression and purification

The human beta-cardiac myosin 15 heptad HMM construct includes residues 1-946 of the *MYH7* gene (GenBank: AAA51837.1) with a leucine zipper GCN4 sequence (MKQLEDKVEELLSKNYHLENEVARLKKLVGER) added after residue 946, followed by a short linker (GSGKL), a C-terminal EGFP tag (M2β 15HPZ.GFP), Avi (GLNDIFEAQKIEWHE) and FLAG (DYKDDDDK) tags. Another construct was generated of just the S1 region which included residues 1-841 of the *MYH7* gene and C-terminal Avi and FLAG tags. The E525K mutation was introduced by Quikchange sited directed mutagenesis into M2β 15HPZ.GFP and M2β S1. The constructs were cloned into the pDual shuttle vector and the initial recombinant adenovirus stock was produced by Vector Biolabs (Malvern, PA) at a titer of 10^8^ plaque forming units per ml (pfu/ml). As previously described, the virus was expanded by infection of Ad293 cells at a multiplicity of infection (MOI) of 3-5 (Swenson et al., 2017). The virus was harvested from the cells and purified by CsCl density sedimentation, giving a final virus titer of 10^10^-10^11^ pfu/ml.

The mouse skeletal muscle derived C2C12 cell line was used to express the cardiac myosin constructs as described previously (Chow et al., 2002; Swenson et al., 2017; Wang et al., 2003; Winkelmann et al., 2015). Briefly, C2C12 cells were grown to ∼90% confluency on tissue culture plates (145/20 mm) in growth media (DMEM with 9% FBS). On the day of infection 20 plates were differentiated by changing the media to contain horse serum (DMEM with 9% horse serum, 1% FBS) and simultaneously infected with virus at 4 × 10^7^ pfu/ml. The cells were harvested for myosin purification 10-12 days after infection and M2β constructs were purified by FLAG affinity chromatography. Actin was purified using acetone powder from rabbit skeletal muscle (Pardee and Spudich, 1982).

### Steady-state ATPase assays

The actin-activated ATPase was examined with the NADH coupled assay as described in previous studies (Rasicci et al., 2021; Swenson et al., 2017; Tang et al., 2021). The ATPase activity was plot as a function of actin concentration and fit to a Michaelis Menten equation to determine the maximum ATPase rate (*k*_cat_) and actin concentration at which ATPase in one-half maximal (*K*_ATPase_).

### In vitro motility assays

The in vitro motility (IVM) assay was performed with M2β 15HPZ.GFP WT and E525K constructs, adapted from previously established protocols (Kron et al., 1991; Rasicci et al., 2021). Briefly, microscope cover slips were coated with 1% nitrocellulose in amyl acetate (Ladd Research) and applied to a microscope slide with double-sided tape to create a flow cell. Myosin in MOPS 20 buffer at concentrations between 72 -90μg/mL (0.4 - 0.5μM) was applied directly to the nitrocellulose surface, and the surface subsequently was blocked with BSA (1 mg/mL). To ensure inactive myosin heads (“dead heads”) were blocked, unlabeled sheared actin (2μM) was added to the flow cell and chased with ATP (2mM). Actin was labeled with phalloidin-Alexa 555 (DsRed filter; excitation/emission 555/588 nm). To initiate motility, an activation buffer containing 0.35% methylcellulose, an ATP regeneration system (2 mM ATP, 5 mg/mL glucose, 46 units/mLpyruvate kinase, and 0.46 mM phosphoenolpyruvate), and oxygen scavengers (0.1 mg/mL glucose oxidase, 0.018 mg/mL catalase) was added to the flow cell. Temperature (25±1°C) was monitored using a thermocouple meter (Stable Systems International). The slide was visualized promptly with a NIKON TE2000 microscope equipped with a 60x/1.4 NA phase objective and a Perfect Focus System. All images were acquired at 1 second intervals for 2 minutes using a shutter controlled CoolSNAP HQ2 cooled CCD digital camera (Photometrics) binned 2×2. Videos were exported to ImageJ and prepared for automated FAST software motility analysis (Aksel et al., 2015), from which >1000 actin filaments from one experiment (i.e. slide) per protein preparation (n=3) at 0.4-0.5μM myosin were compiled for statistical analysis, WT vs. E525K.

### Transient kinetic measurements

An Applied Photophysics stopped-flow equipped with an excitation monochromator, 1.2 ms dead-time, and a 9.3 nm band pass was used for all experiments. The mant fluorescence was examined with 290 nm excitation and a 395 nm long pass emission filter. Single mantATP turnover experiments were performed by incubating M2β constructs (0.25µM) with mantATP (1µM) for 30 seconds and then mixing the complex with saturating ATP (2 mM) and monitoring the fluorescence decay over a 1000 second period. Fluorescence transients were fit to sum of exponentials using the stopped-flow program or GraphPad Prism. For example, the following function was used to fit fluorescence decays, 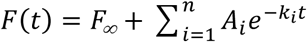 where *F(t)* is the fluorescence as a function of time *t, F_∞_* is the intensity at infinity, *A*_i_ is the amplitude, *k*_*i*_ is the observed rate constant characterized by the *ith* transition, and *n* is the total number of observed transitions.

### Steady-state FRET measurements

Stopped-flow FRET was examined by monitoring the change in donor fluorescence (GFP) using an excitation wavelength of 470 nm and measuring the emission with an interference filter (500-525nm), which eliminated background fluorescence from Cy3ATP. The steady-state FRET efficiency (*E*) was calculated by examining donor quenching using the following equation (Lakowicz, 2006; Tang et al., 2021),

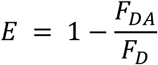

where *F*_DA_ is the donor fluorescence intensity in the presence of acceptor (under conditions of saturating Cy3ATP bound to M2β 15HPZ.GFP), *F*_D_ the donor fluorescence intensity in the absence of acceptor (M2β 15HPZ.GFP bound to ATP). The distance (*r*) between the donor and acceptor was calculated based on the equation below (Lakowicz, 2006),

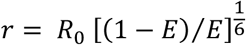

where the Förster distance (*R*_0_), the distance at which energy transfer is 50% efficient, was determined to be 63Å (Fessenden, 2009).

### Time resolved FRET measurements

Fluorescence lifetime measurements of M2β 15HPZ.GFP were performed using time-correlated single photon counting (DeltaPro, Horiba Scientific) with a 479nm pulse diode laser and a 515nm long-pass emission filter. We examined M2β constructs (60 nM) in the absence of nucleotide, presence of 0.1µM Cy3ATP, and in the presence of saturating ATP (1 mM) and 0.1µM Cy3ATP (ATP chase). The fluorescence lifetime decays were fit to two-exponential function and used to determine the average lifetime of the donor alone (τ_*D*_) and donor+acceptor (τ_*DA*_). The FRET efficiency was calculated using the following equation:

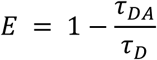

Three technical replicates were performed on a single protein preparation and the FRET efficiency was then used to determine the FRET distance (*r*) as described above.

### Electron Microscopy. Sample preparation and imaging

The samples (WT and E525K) were diluted in 20 mM KCl, 10 mM MOPS, 1 mM EGTA, 0.5 mM ATP, 2 mM MgCl_2_, pH 7.4, to a final concentration of 77 nM and incubated at room temperature for 10 minutes. Negative staining was performed using 400 mesh copper grids with a thin carbon film on top. The grids were glow discharged for 45 seconds at 15 mA in a PELCO easiGlow to make the surface hydrophilic, aiding stain spreading. 5µl of sample was applied to the grid for ∼ 5 seconds and the grid immediately rinsed with 1% (w/v) uranyl acetate for negative staining. Grids were imaged on an FEI Tecnai Spirit Transmission Electron Microscope at 120 kV with a Rio 9 3K x 3K CCD camera (Gatan).

### Calculation of percentage folded molecules

A total of 9 micrographs for each construct were analyzed and visual counting was performed to calculate the percent of folded molecules. The resulting values were used to plot a bar diagram.

### 2-D classification and averaging

A total of 55 micrographs with 2.6 A pixel size were used to extract 1397 (WT) and 1845 particles (E525K) in Relion 3.1.2 (Scheres, 2012). The particles were classified into 50 classes with a mask size of 520 Å and the top 10 classes were used to make the comparison.

### Statistics

For this work, a biological replicate is defined as a new protein preparation, whereas a technical replicate is defined as a repeated experiment from the same stock of protein. Reported sample sizes refer specifically to biological replicates (except where noted), and a minimum of N=3 is reported for all experiments. Groups were allocated as WT vs. E525K, unless otherwise noted, and direct comparisons were made between these two groups. Accordingly, unpaired student’s *t*-tests were performed to compare differences between the groups in all experiments with statistical error. No statistics were performed in the EM data, as there is no error in the open vs. closed 2-D classification. To avoid technical replicates in the in vitro motility assay (i.e. actin filaments from the experiment), sample means of independent experiments were averaged, and the data has been presented as SuperPlots, as previously described (Lord et al., 2020; Rasicci et al., 2021). Stuck filaments (velocity = 0 nm/sec) were excluded from this analysis. Otherwise, no data was excluded from any other reported experiment. The P-values for all statistical comparisons can be found in the supplementary Tables or source files.

## Acknowledgements

This work was supported by NIH grants to HL127699 to CMY, AR072036 and HL139883 to RC, and an AHA Post-doctoral Fellowship to PT.

## Supporting Information

**Table S1.**
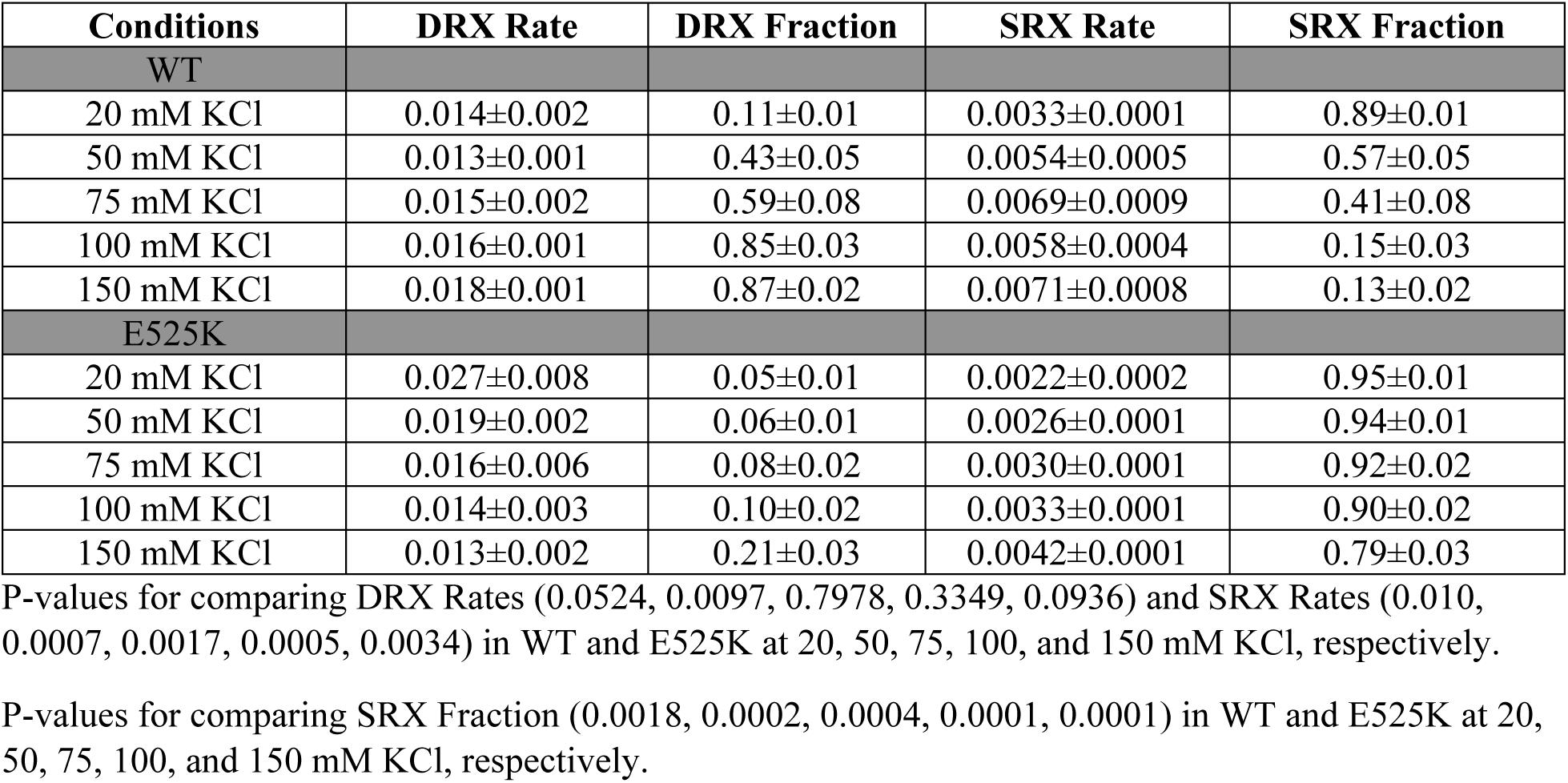
Summary of single mantATP turnover measurements for WT and E525K M2β 15HPZ.GFP (N=3, ±SD).

**Table S2.** Summary of single mantATP turnover measurements for WT and E525K M2β S1 (N=1, ±SE of the fit).

| Conditions | DRX Rate | DRX Fraction | SRX Rate | SRX Fraction |
| --- | --- | --- | --- | --- |
| WT |  |  |  |  |
| 20 mM KCl | 0.037±0.002 | 0.87±0.08 | 0.016±0.006 | 0.08±0.09 |
| 50 mM KCl | 0.033±0.001 | 0.94±0.02 | 0.011±0.004 | 0.03±0.02 |
| 75 mM KCl | 0.031±0.002 | 0.94±0.03 | 0.008±0.002 | 0.03±0.01 |
| 100 mM KCl | 0.029±0.001 | 0.94±0.03 | 0.011±0.004 | 0.04±0.03 |
| 150 mM KCl | 0.025±0.001 | 0.96±0.01 | 0.007±0.002 | 0.03±0.01 |
| E525K |  |  |  |  |
| 20 mM KCl | 0.082±0.003 | 0.70±0.03 | 0.022±0.010 | 0.04±0.03 |
| 50 mM KCl | 0.075±0.002 | 0.85±0.04 | 0.022±0.008 | 0.06±0.04 |
| 75 mM KCl | 0.069±0.002 | 0.88±0.04 | 0.020±0.010 | 0.04±0.04 |
| 100 mM KCl | 0.065±0.001 | 0.88±0.02 | 0.018±0.005 | 0.05±0.02 |
| 150 mM KCl | 0.059±0.001 | 0.91±0.02 | 0.015±0.009 | 0.03±0.02 |

**Table S3.** Summary of kinetic parameters from Cy3ATP binding experiments. (representative data, N=1, ±SE of the fit)

| Parameter | WT | E525K |
| --- | --- | --- |
| $k_{\text{fast}}$ (20 mM KCl) ($\mu\text{M}^{-1}\cdot\text{s}^{-1}$) | 1.9±0.1 | 1.4±0.1 |
| $k_{\text{slow}}$ (20 mM KCl) ($\mu\text{M}^{-1}\cdot\text{s}^{-1}$) | 0.15±0.01 | 0.06±0.01 |
| $A_{\text{fast}}$ (20 mM KCl) | 0.65±0.04 | 0.25±0.05 |
| $k_{\text{on}}$ (150 mM KCl) ($\mu\text{M}^{-1}\cdot\text{s}^{-1}$) | 0.40±0.02 | 0.46±0.03 |

**Table S4.** Summary of steady-state FRET measurements (N=3, ±SD).

| Conditions | FRET Efficiency (%) | Distance ($\text{\AA}$) | Predicted Distance ($\text{\AA}$) |
| --- | --- | --- | --- |
| WT |  |  |  |
| 20 mM KCl | 5.4±0.03 | 101.5±1.0 | 102.0±0.2 |
| 50 mM KCl | 4.8±0.01 | 103.8±0.4 | 110.1±1.7 |
| 75 mM KCl | 3.7±0.06 | 108.4±3.1 | 116.9±3.7 |
| 100 mM KCl | 2.7±0.02 | 114.7±1.5 | 137.9±4.9 |
| 150 mM KCl | 1.7±0.02 | 124.3±2.5 | 141.6±3.0 |
| E525K |  |  |  |
| 20 mM KCl | 5.8±0.08 | 100.3±2.5 | 100.9±0.1 |
| 50 mM KCl | 5.1±0.07 | 102.4±2.5 | 101.0±0.1 |
| 75 mM KCl | 5.5±0.05 | 101.3±1.6 | 101.5±0.3 |
| 100 mM KCl | 4.9±0.01 | 103.1±0.4 | 101.9±0.5 |
| 150 mM KCl | 4.4±0.02 | 105.4±0.9 | 104.2±0.7 |

P-values for comparing FRET Efficiency (0.0013, 0.0018, 0.0001, 0.0001, 0.0001) in WT and E525K at 20, 50, 75, 100, and 150 mM KCl, respectively.

**Table S5.** Summary of time-resolved FRET measurements (N=3, ±SD).

| Conditions | Average Lifetime | FRET Efficiency (%) | Distance ($\text{\AA}$) |
| --- | --- | --- | --- |
| WT |  |  |  |
| GFP-Alone: 20 mM KCl | 2.48±0.01 | | |
| GFP+Cy3ATP: 20 mM KCl | 2.33±0.01 | 6.2±0.3 | 99.1±0.9 |
| GFP+Cy3ATP: 20 mM KCl<br>(ATP chase) | 2.48±0.01 | | |
| GFP-Alone: 150 mM KCl | 2.45±0.01 | | |
| GFP+Cy3ATP: 150 mM KCl | 2.39±0.01 | 2.4±0.3 | 116.8±2.5 |
| GFP+Cy3ATP: 150 mM KCl<br>(ATP chase) | 2.45±0.01 | | |
| E525K |  |  |  |
| GFP-Alone: 20 mM KCl | 2.48±0.02 | | |
| GFP+Cy3ATP: 20 mM KCl | 2.35±0.03 | 5.5±0.9 | 101.2±3.0 |
| GFP+Cy3ATP: 20 mM KCl<br>(ATP chase) | 2.47±0.02 | | |
| GFP-Alone: 150 mM KCl | 2.45±0.01 | | |
| GFP+Cy3ATP: 150 mM KCl | 2.34±0.01 | 4.8±0.8 | 103.7±3.0 |
| GFP+Cy3ATP: 150 mM KCl<br>(ATP chase) | 2.46±0.01 | | |

P-values comparing distance (0.3101, 0.0044) in WT and E525K at 20 and 150 mM KCl, respectively.

Supplementary Movies – In vitro motility of WT and E525K M2β 15HPZ.GFP

**Figure S1.**
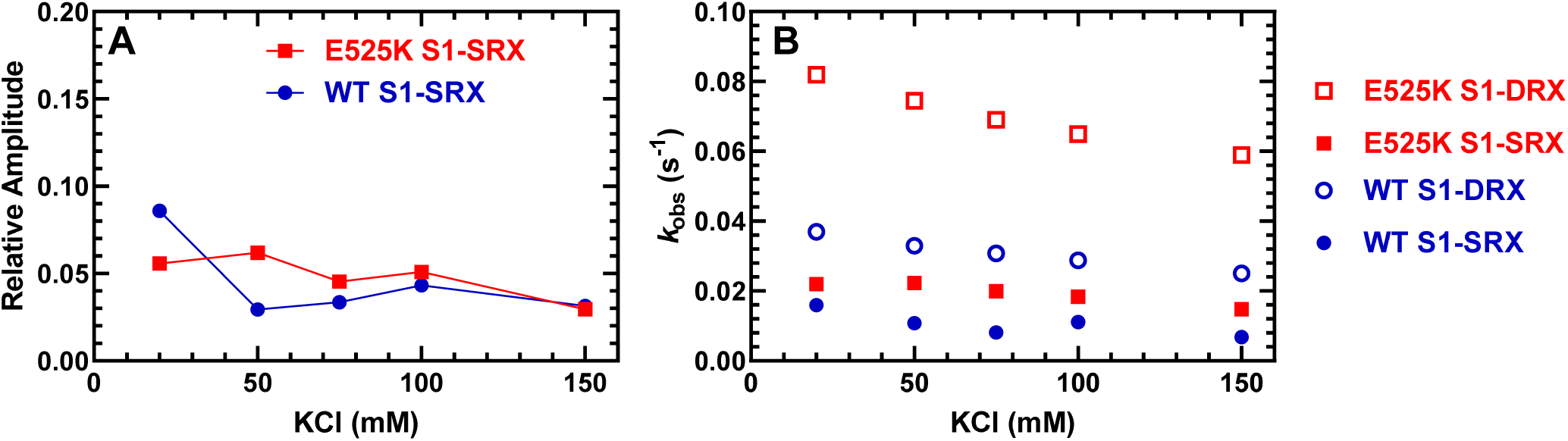
Single turnover with WT and E525K M2β S1 Single ATP turnover measurements with M2β S1. The turnover of mantATP by M2β S1 was examined at varying KCl concentrations as in Figure 3. The mant fluorescence transients were best fit to a three-exponential function. There was a minor very fast phase (∼5% of the signal), which attributed to a small to mantADP release from S1 (∼4 s^-1^) and therefore was not included in the analysis of the other two rate constants which were similar to those observed in Figure 3. A) The relative amplitude of the slow rate constant was used to determine the fraction of heads in the SRX state. B) The slow rate constant (SRX state, circles) was similar in WT and E525K while the predominant rate constant (DRX state, squares), which dominated the fluorescence transients (90-95% of the signal), was 2-fold higher in E525K compared to WT. All rate constants were relatively insensitive to KCl concentration (see Table S2 for summary of values).

**Figure S2.**
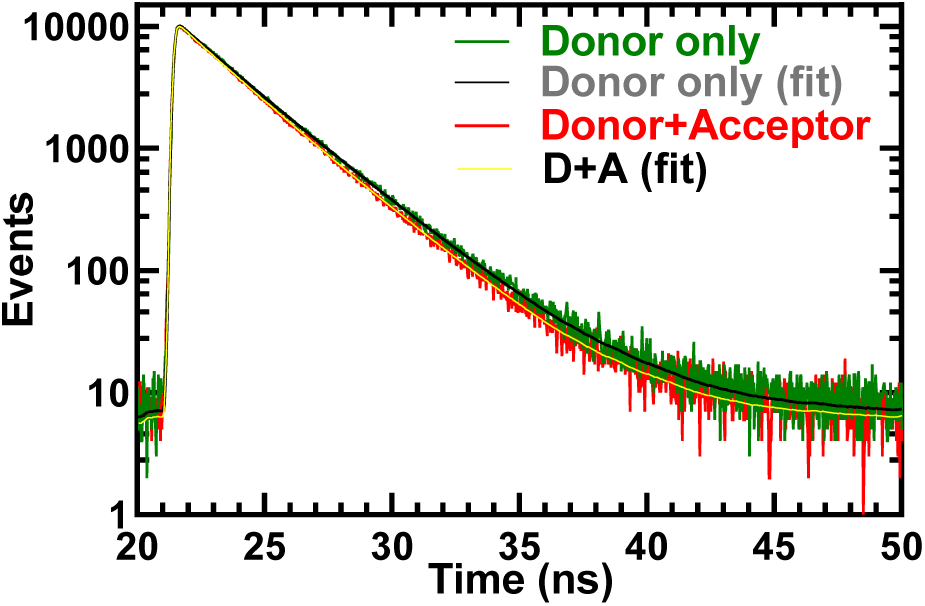
TR-FRET decays Representative fluorescence lifetime decays. Fluorescence lifetime decays were collected by exciting the donor (GFP) using a 479 nm picosecond laser and examining donor fluorescence emission (515 long-pass filter) in the presence and absence of acceptor (Cy3ATP). Time correlated single photon counting was used to collect lifetime decays. The donor only (60 nM M2β 15HPZ.GFP) is compared to the donor+acceptor (60 nM M2β 15HPZ.GFP+0.1µM Cy3ATP) in 20 mM KCl conditions. The decays were fit well to a two-exponential function to determine the average lifetime (Table S5 for summary of time-resolved FRET).

**Figure S3.**
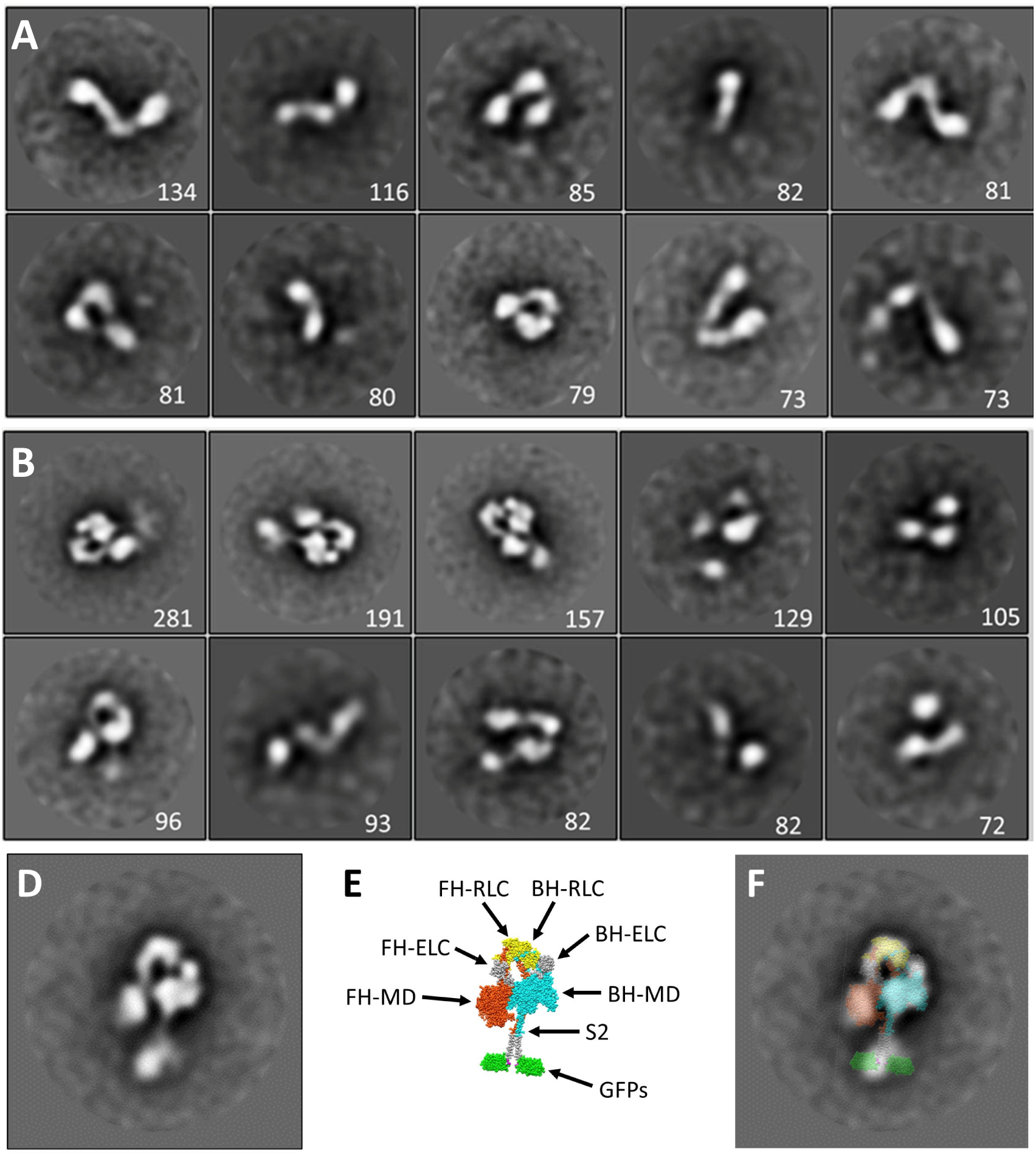
EM raw data and image processing 2D class averaging of M2β 15HPZ.GFP negative stain images. (A) WT and (B) E525K. The numeral in each panel indicates the number of particles contributing to the particular class average. (D)-(F) Superposition of M2β 15HPZ.GFP atomic model on E525K class average. (D) Class average. (E) Atomic model based on 5tby (Fig. 1). (F) Superposition, demonstrating good agreement between EM imaging and model, supporting our identification of the molecules as *bona fide* IHMs.

**Figure S4.**
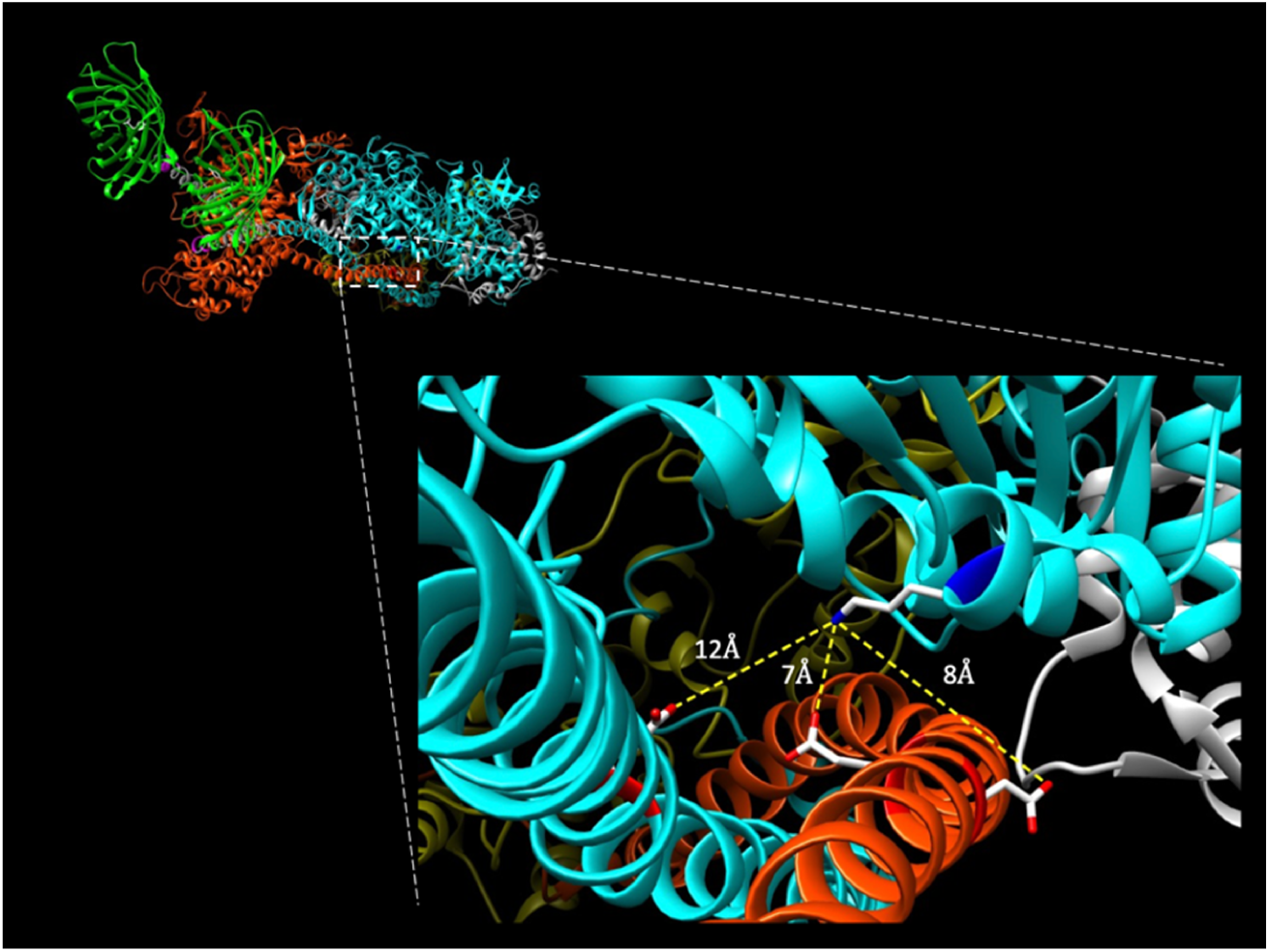
Potential intramolecular interactions of E525K with S2 Potential intramolecular interactions of E525K with S2. *Top left*. Oblique view of folded myosin hexamer from the C-terminal end. Note C-terminal GFP molecules for orientation of molecule with blocked head (BH, cyan) and free head (FH, orange) of myosin. *Inset*. Potential intramolecular interactions between the mutated 525K residue (blue, positively charged amine group) on BH of myosin with nearby acidic residues on S2 (red, negatively charged carboxylic acids) of both BH and FH. Highlighted residues include E902 on S2 of BH and E894 and D896 on S2 of FH. Intramolecular interactions were mapped using UCSF Chimera on same closed structure (5TBY template) described in Figure 1 (Alamo et al., 2017; Pettersen et al., 2004).

